# Massive loss of transcription factors and the initial diversification of placental mammals

**DOI:** 10.1101/2021.03.26.436744

**Authors:** Xin-Wei Zhao, Jiaqi Wu, Hirohisa Kishino

**Author notes:** Correspondence to (X.-W.Z.). These authors contributed equally to this work (Xin-Wei Zhao, Jiaqi Wu).

## Abstract

As one of the most successful categories of organisms, mammals occupy a variety of niches on earth as a result of macroevolution. Transcription factors (TFs), the basic regulators of gene expression, may also evolve during mammalian phenotypic diversification and macroevolution. To examine the relationship between TFs and mammalian macroevolution, we analyzed 140,821 *de novo*-identified TFs and their birth and death histories from 96 mammalian species. Gene tree vs. species tree reconciliation revealed that mammals experienced an upsurge in TF losses around 100 million years ago and also near the K–Pg boundary, thus implying a relationship with the divergence of placental animals. From approximately 100 million years ago to the present, losses dominated TF events without a significant change in TF gains. To quantify the effects of this TF pruning on mammalian macroevolution, we analyzed rates of molecular evolution and expression profiles of regulated target genes. Surprisingly, TF loss decelerated, rather than accelerated, molecular evolutionary rates of their target genes, suggesting increased functional constraints. Furthermore, an association study revealed that massive TF losses are significantly positively correlated with solitary behavior, nocturnality, reproductive-seasonality and insectivory life history traits, possibly through rewiring of regulatory networks.

## INTRODUCTION

Gene expression patterns vary among species—even closely related taxa that share highly similar genomic sequences. These differences in gene expression and regulation are believed to be the major source of species phenotypic variation as well as important factors in evolution (1). Transcription factors (TFs), which are essential gene regulators, perform the initial step of DNA decoding (2).

TFs have many important functions in eukaryotes (3,4). As revealed by laboratory experiments, the ability of TFs to drive phenotypic changes has long been known. For example, HOX TF genes play a key role in proper body pattern formation (5), while SRY, a TF gene, is important for sex determination (6,7). In addition, mutations of TF genes have many unexpected consequences, such as the formation of induced pluripotent stem cells (8) and cancer (9,10).

Simplification and complication are both critical aspects of macroevolution. Simplification, that is, the reduction of biological complexity to varying degrees, has received less scientific attention than complexity. Examples of simplification-driven diversification across the tree of life include simplification events during the early history of metazoans, convergent losses of complexity in fungi, and simplification during early eukaryotic evolution [reviewed by (11)]. Nonadaptive simplification, such as drift, can lead to the accumulation of slightly deleterious mutations in bacteria (12). Adaptive genome reduction may also explain some important stages of eukaryotic evolution, such as the simplification of animal metabolism (13). The ‘less-is-more principle’ suggests that loss of gene function is a common evolutionary response of populations undergoing an environmental shift and, consequently, a change in the pattern of selective pressures (14). In this regard, TF patterns may naturally evolve along with organismal macroevolution.

The important components of metazoan and embryonic-plant TF kits were present even earlier in their respective single-cell ancestors (15). Given that the origin and expansion of TFs occurred long before the big bang of speciation, macroevolution has likely been driven by a more direct factor, possibly TF loss. TF loss leading to major diversification has occurred in eukaryotes–for instance, the convergent simplification of adaptin complexes in flagellar apparatus diversification (11). As another example, the unexpectedly complex list of Wnt family signaling factors evolved in early multicellular animals about 650 million years ago (Mya) (16). Functional and phenotypic diversification of the mouth was caused by the loss of Wnt family signaling factors during animal evolution (17).

How TFs work when mammalian species quickly adapt to changing environments via macroevolution is poorly understood. Advances in comparative genomics have clearly shown that the exclusive use of genes as evolutionary units is an oversimplification of actual evolutionary relationships (18,19). The related concept of orthologous groups refers to a set of homologous genes that evolved from a single ancestral gene after a given speciation event. Given the close connection between orthologous groups and evolutionary events, we used orthologous groups, rather than genes or gene families, as the basic unit in this study to detect gain and loss events of global TFs in mammalian evolutionary history. Because of the lack of phenotypic data related to orthologous groups and the high resolution of orthologous genes in loss events, we used orthologous TFs to identify the association between TF loss and traits. Here, we show the pattern of TF loss enrichment in the macroevolutionary process.

The role of TFs in macroevolutionary processes is further discussed by describing the correlation between TF loss, target gene (TG) expression and molecular evolutionary rate, and also between TF loss and species traits. The results of our analysis may provide new insights into the role of TF loss under various macroevolutionary models as well as its contribution to the rapid adaptation of species to different environments.

## MATERIALS AND METHODS

### Mammalian TFs

Based on our previous research, we obtained 140,821 TF proteins, nearly all of which were mammalian TF proteins (20). To assess the accuracy of TF annotations, we compared our list of 1625 human TFs (extracted from the 40,821 TFs in 96 mammalian genomes shown in Table S1) with two well-known TF databases, AnimalTFDB3 (21) and HumanTFs (4) (Fig. S1). The number of human TFs in these two databases is similar to that of our list: 1639 in humanTFs and 1665 in AnimalTFDB3. A total of 1402 TFs are listed in all three databases, while 82 are unique to HumanTFs, 123 are found only in AnimalTFDB3, and 140 are restricted to our human TF list. All three databases are based on similar DBD and HMMER pipelines.

AnimalTFDB3 contains 125,135 TFs from 97 genomes ranging from *Caenorhabditis elegans* to mammalian species such as humans, whereas our database contains 140,821 TFs and focuses on 96 mammalian species. The latter database may thus provide better insights into the evolutionary history of TFs in mammals. To avoid the limitations of mammalian species, we further annotated our database with information on orthologous groups from OrthoDB. In this way, we can trace TFs in mammalian species back to those of bacterial species.

To further confirm events occurring in the common ancestor of mammals, we conducted a hidden Markov model (HMM) search on 11 outgroup species (four birds, four reptiles, two amphibians and one coelacanth) and collected all genes predicted to encode TFs.

### Multiple alignment of mammalian orthologs

We downloaded 96 complete mammalian genomes from GenBank and used a custom Perl script to extract protein-coding sequences of each species. A gene pool of 21,350 mammalian genes was constructed based on NCBI genomic annotations. Using the results of the HMM search and annotations in OrthoDB, we assigned TFs to 1651 mammalian orthologous groups. We confirmed the homology of outgroup-TFs and mammal-TFs by a local blast search [blastp,(22)] and assigned outgroup-TFs to mammalian orthologous groups.

We generated a multi-sequence file of all genes in the gene pool for each of the orthologous groups. Codon-level alignments were performed in Prank v.170427 (23). Sites with less than 70% coverage across all species, as well as sequences with less than 30% coverage among gene loci, were removed from the alignment.

### Inference of gene trees

We estimated the maximum likelihood tree for each gene using IQ-TREE software (24), which automatically performed model selection and determined the best data partitions. The best evolutionary model for each gene was independently selected based on the Bayesian information criterion and used for inference of the nucleotide tree. All gene trees were calculated using 1,000 bootstrap replicates.

### Species tree inference and divergence time estimation

We selected 823 genes that were single-copy in mammals according to Wu et al. (25) and present in all 96 mammals. The 823 genes were used to infer a species tree using the coalescent-based Njst method (26). The topology of the inferred species tree was consistent with that of Tarver et al. (2016), who placed treeshrew (*Tupaia chinensis*) as the root lineage of Glires (27). The phylogenetic position of treeshrew is not yet resolved, however, and several researchers consider treeshrew to be the root lineage of Euarchonta, not Glires (28). As an alternative species tree in this study, we consequently used a tree in which the position of treeshrew was fixed at the root of Euarchonta rather than Glires. Following the same method as Wu et al. 2017, we estimated the divergence times of 96 mammals based on the inferred branch effect (genomic rate × genomic time) and fossil calibrations (25).

### Reconstructing the history of TF duplication and loss

A history of gene duplication and loss was estimated by reconciling gene trees with the species tree. To focus on these events during mammalian evolution, TFs were categorized into orthologous groups (OrthoDB, https://www.orthodb.org/), each having a single ancestor at the root of mammals. Out of 140,821 TFs from 96 mammalian species, OrthoDB included nearly 120,000 TFs from 82 species. These TFs were assigned to 2,880 orthologous groups. A phylogenetic tree was constructed for each of the 1,651 orthologous groups containing more than three members. To infer the history of TF duplication and loss, each of the 1,651 phylogenetic trees was reconciled with the species tree according to the criterion of parsimony using NOTUNG 2.9 (29). To take into account inconsistency due to incomplete lineage sorting, we established a branch length threshold in terms of *N_μ_*. Branches with lengths under the threshold were treated as weak branches and rearranged by NOTUNG. For each orthologous group tree, the value of μ corresponded to the average path length from root to terminals divided by the calculated age of the root. We assumed an average generation length of 10 years and considered four possible values for effective population size, namely, 10^4^, 10^5^, 10^6^ and 10^7^, and three tree topologies. The events of orthologous groups were aggregated to obtain the numbers of births and losses of TFs along each branch of the species tree. In this paper, we present the results of the analysis based on an effective population size of 10^6^. Other effective population size threshold settings gave similar results. The sequence depth of platypus was relatively low (6×). To confirm that the inclusion of this species did not lead to biased results, we downloaded 11 more outgroups and 82 more mammalian species, including one fish, two amphibians, four reptiles, and four birds, for use in tree construction. Using the 11 outgroups, 982 TF orthologous groups were traced and analyzed by the above-mentioned method. Divergence times of outgroups were taken from the Timetree (30) database.

### The effect of TF loss on TG molecular evolutionary rates

To examine long-term effects, we compared rates along the terminal branches of a TG tree between species that possess the TF and species that do not. Rates of molecular evolution were estimated by comparing the split decomposition of gene trees with that of the species time tree as follows. After fixing gene tree topologies to be consistent with the species tree, we estimated branch lengths of all 21,350 genes using PAML v.4.9 (31). Because many of the genes had missing taxa, we designed a split decomposition-based method to align gene-tree branches. In this approach, each tree was defined by a unique set of splits that could explain the topology of the gene tree. We assigned the splits so that the same split in the species tree encompassed the same branch and then generated a branch length table for 21,350 gene trees. For branches on which a loss event occurred, only the exact same split type on both the TG branch and the corresponding species-tree branch was considered. Rates of molecular evolution were estimated as the ratios of branch lengths to the relevant time periods in the species tree.

### The effect of TF loss on TG expression profiles: human–mouse comparison

Gene expression data of 15,796 orthologous human and mouse genes in five organs (cerebellum, heart, kidney, liver and testis) were retrieved (32). The average levels of these expression data, which were standardized as transcripts per million kilobases, were similar between humans and mice in these organs. Regulatory information on TFs and their TGs was obtained from TRRUST v.2 (33). To analyze the effect of TF losses on the expressions of their TGs, we compared expression profiles between three types of genes: (1) mouse orthologs of the TGs of human TFs without orthologs in mice, (2) human orthologs of TGs of mouse TFs without orthologs in humans and (3) TGs of TFs orthologous between humans and mice.

### Association study of ecotype traits and TF presence/absence

We manually collected the ecotype traits of 96 mammalian species from the Animal Diversity Web (http://animaldiversity.org/). The presence and absence of TF orthologs in each species was recorded as 1 and 0, respectively, based on mammal-terminal single-copy orthologs. The ecotypes were binary and were recorded as 1 and 0. We identified significant associations using the lasso logistic regression procedure as implemented in the glmnet package in R.

## RESULTS

### TFs were simplified during mammalian evolution

The most recent common ancestor of extant mammals dates back to approximately 170 Mya (Fig. S2). After splitting from Marsupialia, Placentalia diverged into Afrotheria, Xenarthra and Boreoeutheria about 100 Mya. To analyze the history of TF birth and death events, we *de novo* identified 140,821 TFs from 96 mammalian species and grouped them into 1,651 orthologous groups (Fig. S3), each having a single origin at the root of mammals (Table S1). The history of TF duplication and loss in each orthologous group was estimated by reconciling the gene tree with the species tree (Fig. S4). We applied four levels of effective population size—10^4^, 10^5^, 10^6^ and 10^7^—as a threshold when considering incomplete lineage sorting. Although the number of events varied extensively among the four population-size thresholds, the overall trend was quite similar in regard to TF events (Fig. S5). As a representative example of the overall trend, but not exact numbers, the results obtained using an effective population size of 10^6^ are shown in Fig. 1.

**Figure 1.**
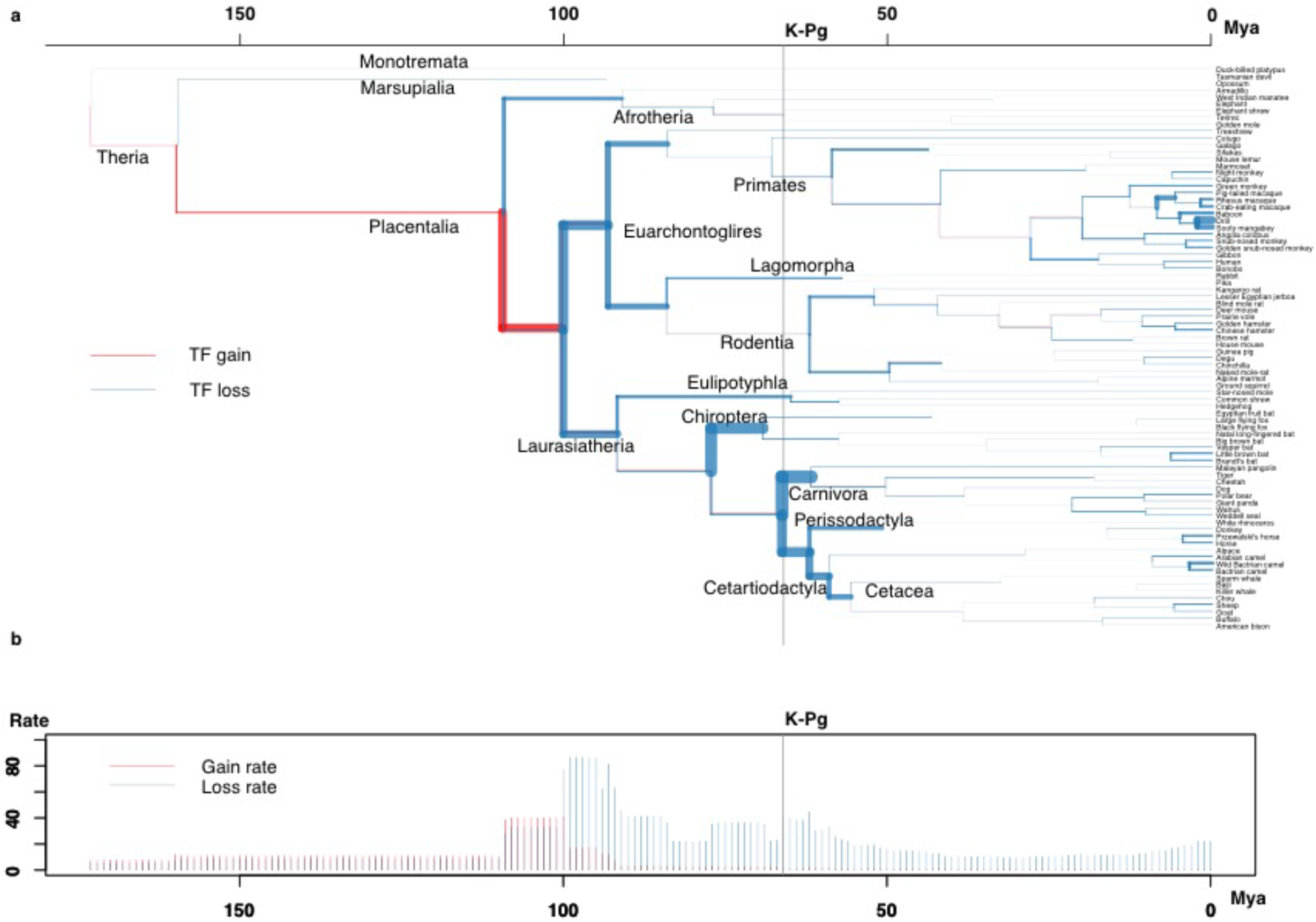
Simplification of ancient transcription factors (TFs) during mammalian evolution. (A) Atlas of TF gain and loss of 982 orthologous groups during mammalian history. The width of each branch is proportional to the rate of TF gain (or loss). Time scale is million years ago (Mya).

To avoid noise arising from the platypus data, we analyzed an additional 11 outgroups and 82 mammalian species (Figs. 1 and S6). When the effective population size was set to 10^6^ in our analysis, the rate of TF amplification before the divergence of Placentalia and Marsupialia was slightly higher than the rate of loss. From approximately 170 to 100 Mya, the pattern of TF gain and loss varied among all mammal TFs (Fig. S2b) and ancient mammal TFs (Fig. 1) but then was identical. This variation may be due to the poor quality of the platypus data. Most TFs arose before the common ancestor of placental animals. TFs were rapidly lost, however, during the evolution of placental mammals, whereas only a few new TFs were generated.

Early Euarchontoglires, Laurasiatheria and marsupials appeared between 100 and 95 Mya and underwent initial diversification. During this time, the average TF loss rate reached its first peak of 90.7/My (Fig. 1b). We repeated the analysis by successively assuming Atlantogenata, Exafroplacentalia and Epitheria topologies at the same threshold level. The events detected under the three different topological arrangements were very similar (Fig. S7). The second peak of TF loss occurred near the Cretaceous–Paleogene (K–Pg) boundary, at 66 Mya. The TF loss rate was as high as 49.4/My within a 4-Mya window of the K–Pg boundary.

As shown in Fig. 1b, the most rapid TF gains mainly occurred during the early stages of mammalian history; this was especially the case for Boreoeutheria, with 1,462 TF gain events at corresponding rates of 159.5/My. After this period, TF loss dominated. We detected 1,287 TF loss events occurring at a rate of 186.3/My along the ancestral branch of Euarchontoglires. In addition, 1,211 TF loss events at a rate of 145.1/My were inferred along the ancestral branch of the Laurasiatherian lineage, while 1,482 taking place at a rate of 81.2/My were uncovered on the ancestral branch of marsupials. TF loss also occurred rapidly on subsequent branches, especially where common ancestors of different mammalian orders started to differentiate, consistent with the above results. For example, estimated loss rates on Chiroptera, Perissodactyla and Carnivora ancestral branches were 208.9, 84.4 and 236.4 /My, respectively. During this time period, the number of species increased significantly. Highly diversified lineages would be expected to contain more highly diversified genes as well, but we found the opposite to be true. The TF pool became simplified during mammalian species diversification, thus indicating that such an event may have contributed to the early diversification of mammals. In contrast, gain events did not follow any specific pattern; therefore, the amplification of TFs may have contributed little to mammalian species diversification.

### The effect of TF loss on relevant regulatory systems

To quantify the effect of TF loss on TF TGs [TF-TG data from TRRUST (33)], we examined TG evolutionary rates. Figure 2, which compares the evolutionary rates of 1-to-1 orthologous TGs with TFs and the evolutionary rates of TGs without TFs, clearly shows the decelerating effect of TF loss on TGs over the long term (Table S2). Because the rate of molecular evolution of a gene is negatively correlated with the strength of a functional constraint, the loss of a TF is expected to have a role in the adaptive evolution of the regulatory system (34).

**Figure 2.**
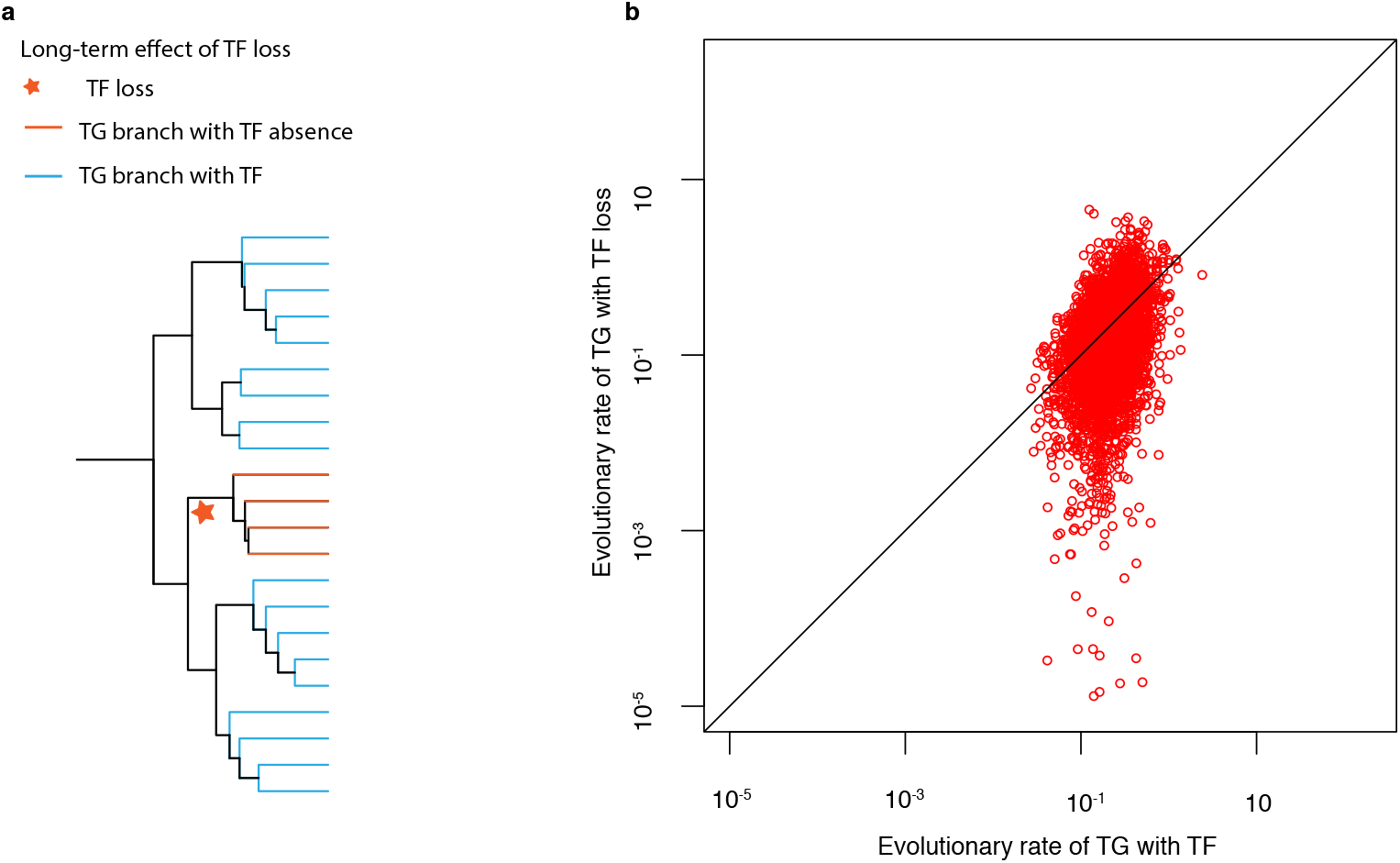
Deceleratory effect of the loss of transcription factors (TFs) on molecular evolutionary rates of their target genes (TGs). (A) Comparison of the evolutionary rates of TGs in species in the subtree of the TF event (colored in red) with the rates of TGs in other species. (B) Long-term effect of TF loss on the evolutionary rate of TGs. X-axis: evolutionary rate of a TG still controlled by a TF; y-axis: evolutionary rate of a TG whose TF has been lost.

The molecular evolutionary rate of a gene is negatively correlated with its expression level (35). To discern the effect of TF losses on expression profiles of their TGs, we compared TG expression profiles between humans and mice. To avoid noise, we used TGs actively regulated by TFs from the TRRUST database. Among these TFs, 614 were present in both humans and mice. In addition, 54 TFs were present in humans but had been lost in mice, while 34 TFs were found in mice but not in humans. In humans and mice, TGs lacking TFs had higher expression levels (Fig. 3). High expression levels can promote a reduction in the rate of gene evolution. This observation is also consistent with our earlier finding that TF loss leads to a decline in TG evolutionary rates. Functional annotation of TGs revealed functions mainly related to the cell cycle, cell migration, signal transduction, and inflammation. High levels of cyclin and signal transduction protein expression suggest that mouse organs undergo higher rates of chemical reactions, and cell migration and inflammation-related proteins play important roles in immunity. These compounds are also related to the small body size, higher somatic mutation rate and shorter life cycle of mice, which results in rapid generational changes and environmental adaptation.

**Figure 3.**
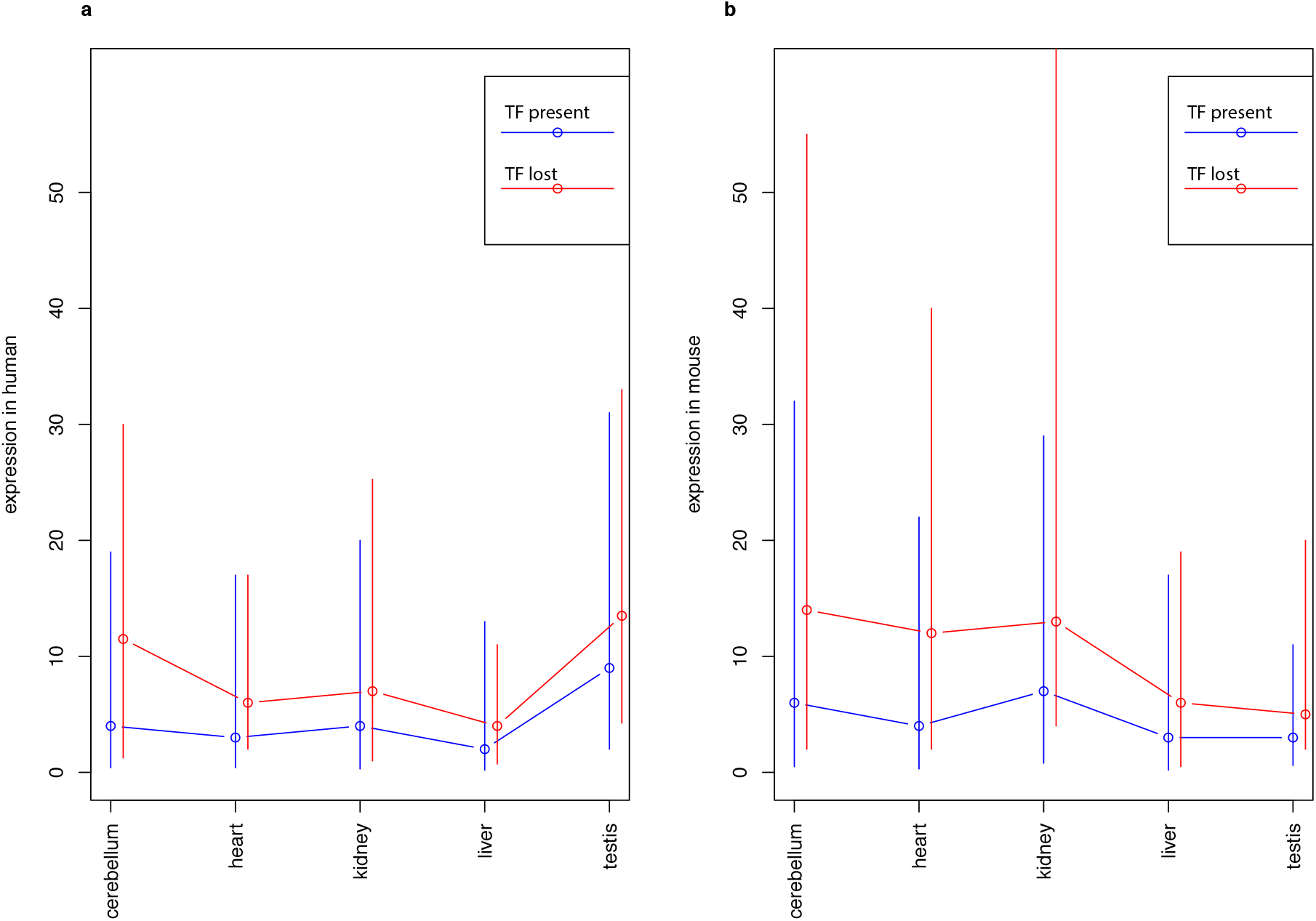
Expression profiles of TGs of TFs present or absent in humans and mice. (A–B) Expression profiles of human (A) and mouse (B) TGs. The circles are median values. The vertical lines are interquartile ranges.

### Association with trait values

To examine the effect of TF loss on life history traits, we regressed four binary traits—sociality, diurnality, reproductive seasonality, and insectivory—on the states (presence/absence) of 1-to-1 orthologous TFs via lasso logistic regression (Fig. 4, Fig. S8, Table S3).

**Figure 4.**
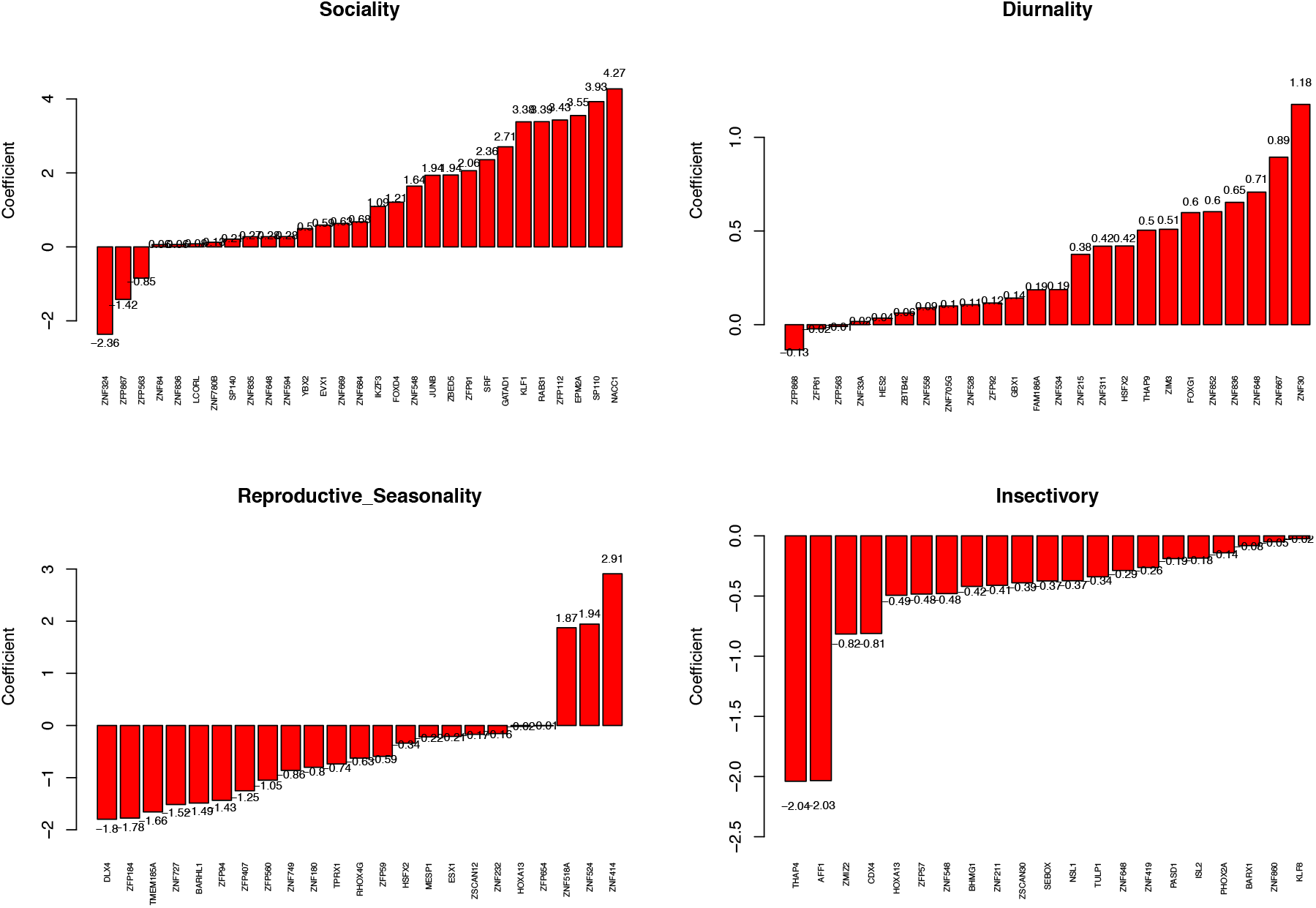
Association of TF loss with mammalian traits. The x-axis shows TF genes significantly related to the indicated trait, and y-axis values are lasso logistic regression coefficients pre-screened with a chi-square test (*P* < 0.05).

TFs, especially members of the zinc-finger protein with Krüppel-associated box domain (KRAB-ZNF) family, are prominent candidates for a role in mammalian speciation (36). Consistent with this idea, ZNFs or KRAB-ZNFs comprised half of the TFs significantly associated with the four life history traits (Table 1).

**Table 1.**
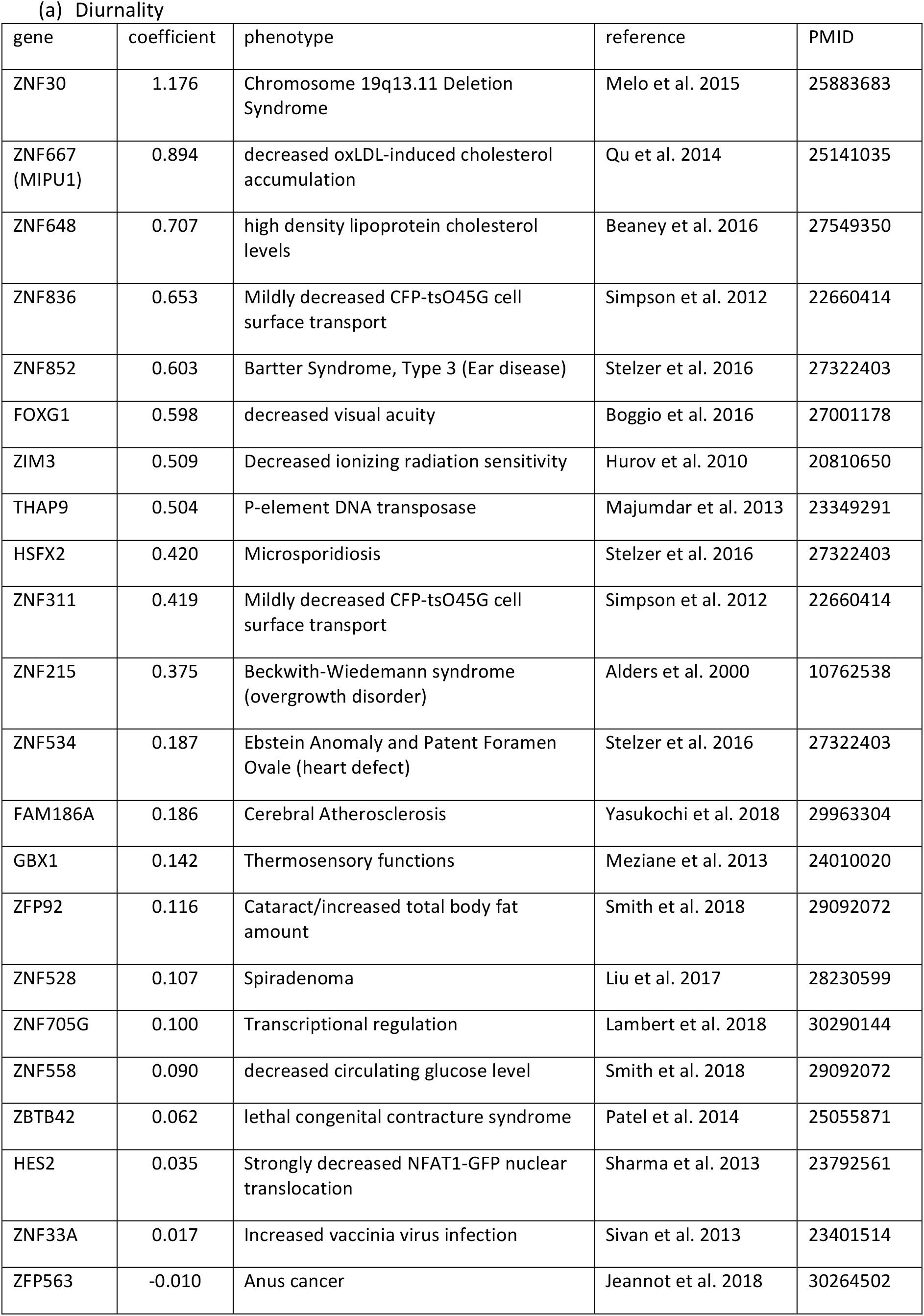
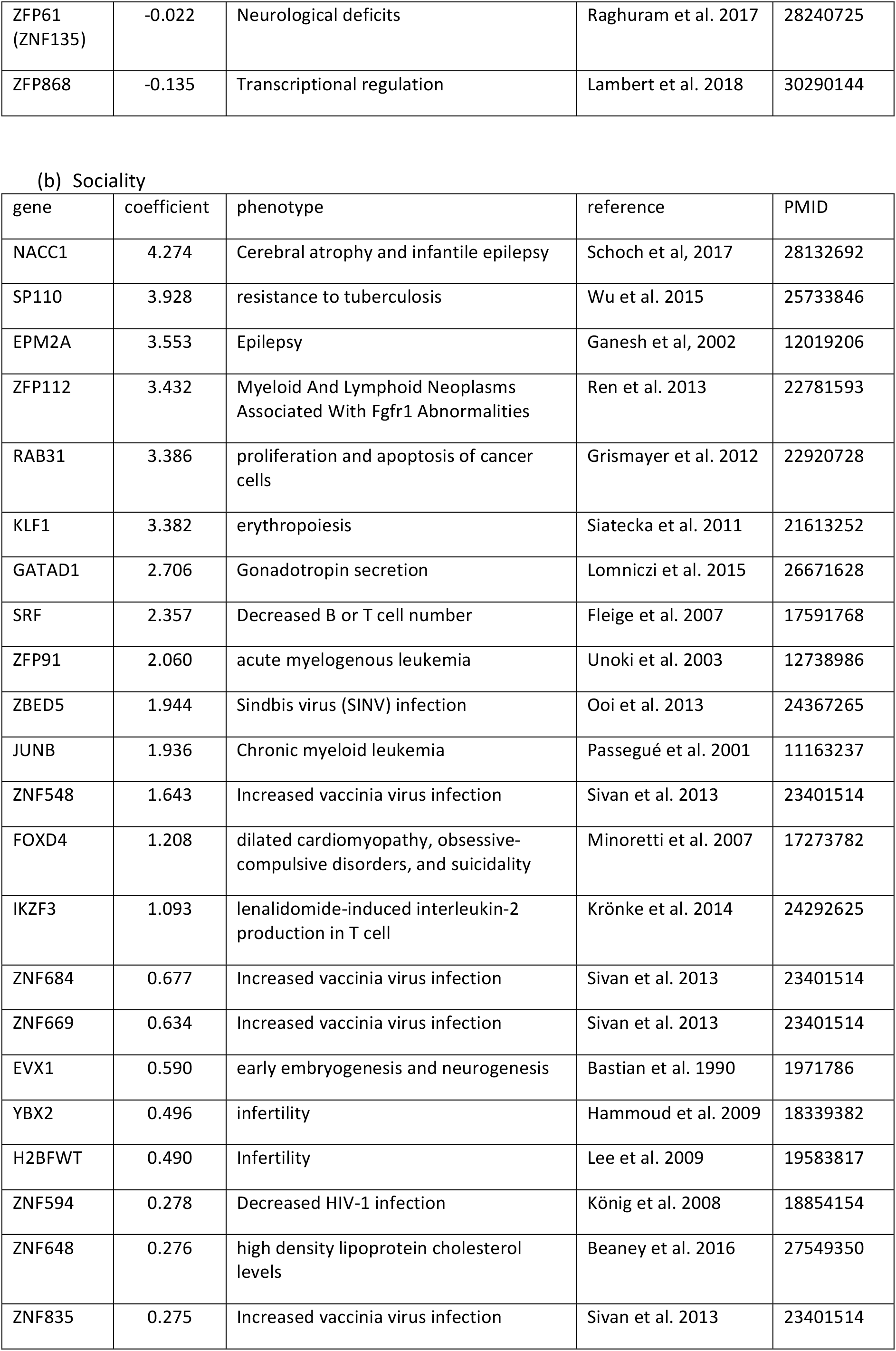
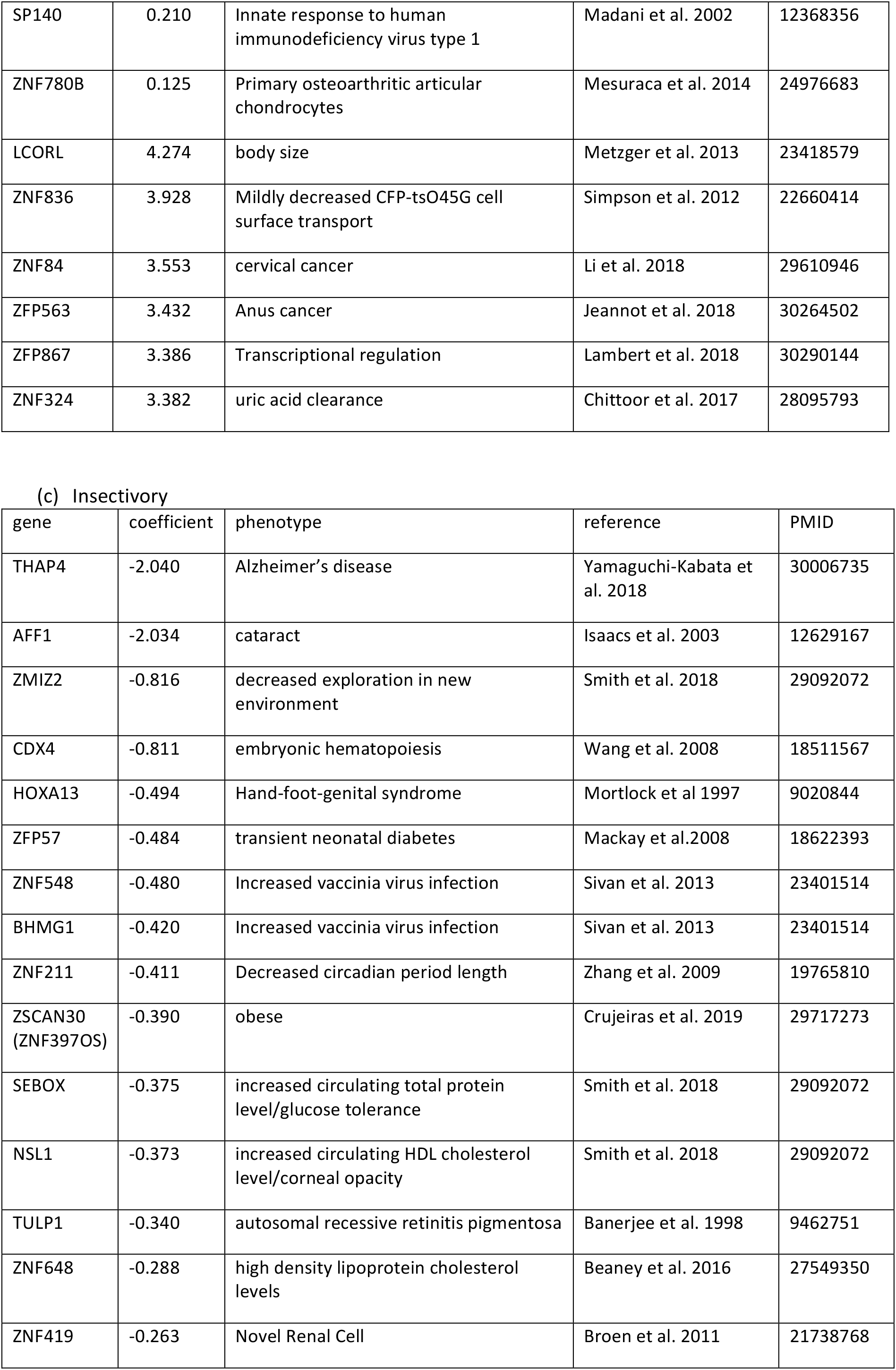
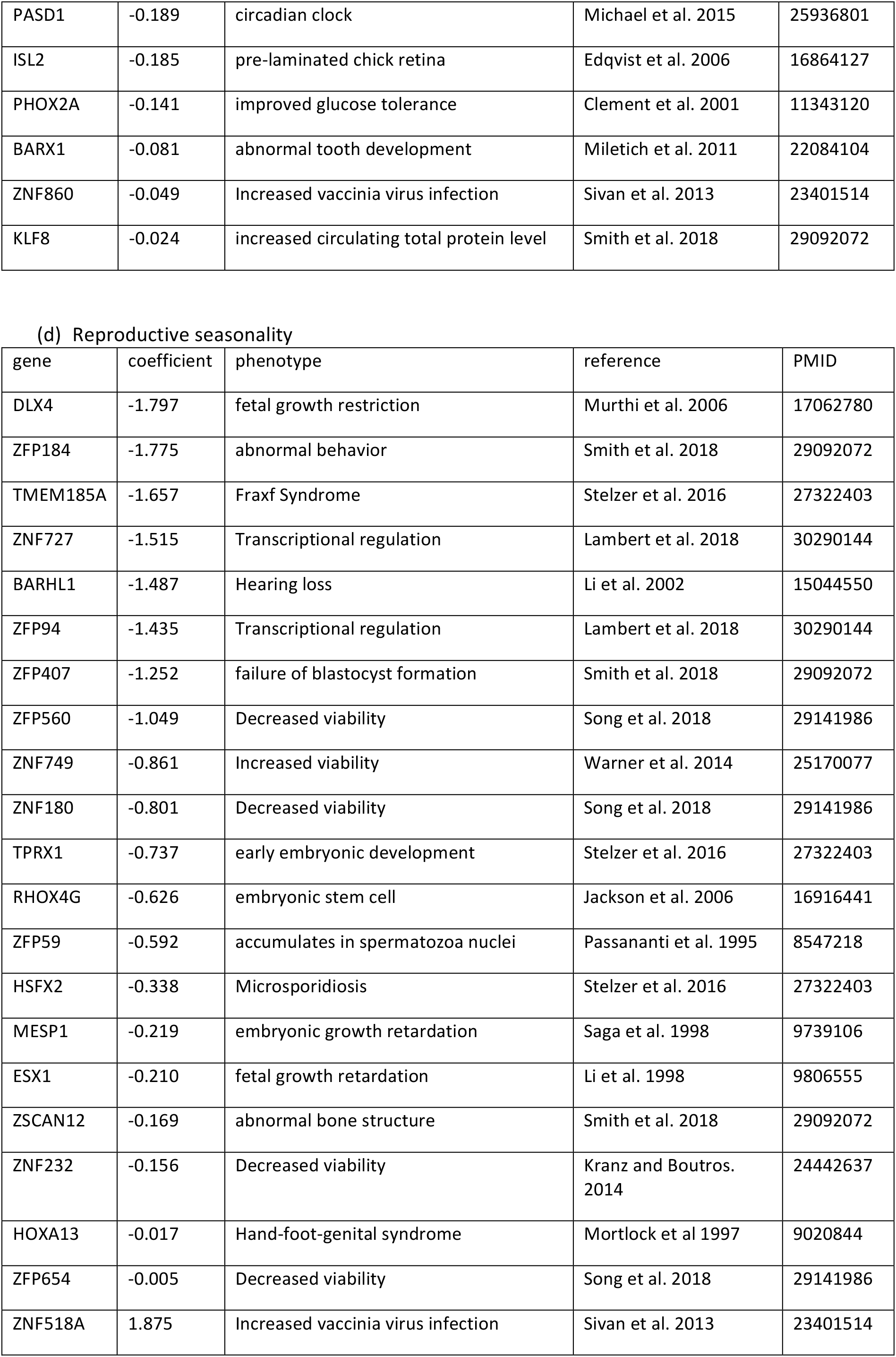
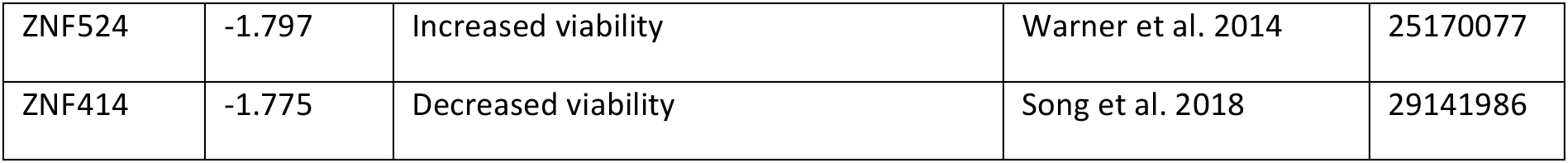
Phenotypes of TF genes with significant trait associations.

In total, 24 TF genes were significantly associated, mostly positively, with diurnality. In other words, TF loss during mammalian evolution was favorable to nocturnal life. The ancestral trait in mammals before the K–Pg boundary was likely nocturnal (25); mammalian species that are still nocturnal have lost these TFs.

Starvation and a cold environment have caused the transition from nocturnal to diurnal habits in mammals, and metabolic balance has been identified as a potential common factor affecting circadian rhythm organization (37). Functions of TFs associated with diurnality shed light on their adaptation. Overexpressed ZNF667 (MIPU1) decreases oxLDL-induced cholesterol accumulation (38). ZNF648 is associated with high-density lipoprotein cholesterol levels *(39).* Knockout of the ZFP92 gene increases the amount of total body fat (40). GBX1-knockout mice exhibit reduced thermosensory functions (41). Changes in the metabolic balance of fat might affect changes in tolerance to starvation and cold temperature. A higher hunger and cold tolerance is likely to promote adaptation to a nocturnal situation.

As for sociality, which exhibited a trend similar to that of diurnality, 26 of the 29 associated TF genes had positive correlation coefficient values. This result indicates that the presence of these significantly related TF genes is positively correlated with social traits; that is, the loss of these TFs is correlated with the solitary trait.

Highly social species have an increased risk of parasitic infection, which necessitates a heavy investment in immune functions (42). Among genes associated with sociality, half were related to immunity-associated phenotypes (Table 1). Sociality has different effects on shape and behavior, including the relative size of the brain and the prevalence of infanticide [reviewed by Silk et al. (43)]. *De novo* mutations in NACC1 can lead to cerebral atrophy and infantile epilepsy (44). Epilepsy can have significant negative consequences on a child’s social development (45). EPM2A can also lead to epilepsy (46). The ancestor of placental mammals was likely solitary (25). Consequently, TF loss helps preserve sociality, and animals without TF loss switched to a solitary lifestyle.

TF genes significantly related to insectivory showed an obvious trend, and their coefficient values tended to be negative. This result indicates that mammalian species that retained the insectivory trait lost these TFs.

The type of food ingested by mammals is also evolutionarily very important. Among genes associated with the insectivory trait, the knockout of SEBOX, PHOX2A and KLF8 genes in mice has been found to increase levels of circulating total protein or improve glucose tolerance (40,47). Knockout of NSL1 and ZNF648 leads to increased circulating cholesterol levels (39,40). According to various knockout data (40,48), AFF1, NSL1 and TULP1 are related to vision. Insectivorous mammalian species usually have weak eyes. TF loss helps preserve insectivory, and animals without TF loss became non-insectivorous.

In terms of reproductive seasonality, coefficient values of 20 of the 23 TFs significantly associated with this trait were all negative. Similar to the trend exhibited by the insectivory trait, this result indicates that TFs positively correlated with the reproductive-seasonality trait have been lost. More than half of the reproductive seasonality-associated TFs were ZNFs. KRAB-ZNFs play a major role in the recognition or transcriptional silencing of transposable elements, and transposon-mediated rewiring of gene regulatory networks has contributed to the evolution of pregnancy in mammals (49). With regard to the phenotype associated with these genes (Table 1), 13 out of 23 are related to fetal viability or formation. Seasonally breeding mammals with these TF losses most likely have better survival chances. TF loss helps preserve reproductive seasonality; animals without TF loss have switched to yearly reproduction. The loss of TFs may therefore have bolstered the inheritance of the ancestral trait of seasonal reproduction (25) in the affected species.

## DISCUSSION

As shown in Fig. 1b, the three branches of Afrotheria, Euarchontoglires and Laurasiatheria experienced a large number of TF loss events approximately 100 Mya, which corresponds to the first peak of TF loss in mammalian evolutionary history. The likely place of origin or current extant range of Afrotheria, Euarchontoglires and Laurasiatheria clades is Africa, Europe and the northern supercontinent of Laurasia, respectively (50,51). This distribution indicates that the living environment of these organisms has likely changed compared with their common ancestor; in other words, their niches and eco-traits may have changed. In contrast, a significant correlation exists between traits and the loss of TFs (Fig. 4). This correlation suggests that TF loss is beneficial to the survival of species undergoing niche and environmental changes and can therefore become fixed in those that survive. At the second peak of TF loss, which took place near the K–Pg boundary, clusters of loss events are apparent on branches, especially the three branches leading to Carnivora, Perissodactyla and Cetartiodactyla. Members of Carnivora are basically carnivores, while Perissodactyla comprises mainly herbivores, and Cetartiodactyla contains both types of feeders. The feeding preferences of the three taxa have thus clearly and rapidly diverged from those of their common ancestor, and the massive loss of TFs in this transition may also have been beneficial to species survival. This phenomenon suggests that TF loss plays an important role in species adaptation to environmental change and macroevolution.

The loss of genes is generally thought to lead to non-functionalization (52). These genes are usually considered to be redundant parts that can be replaced by other genes or provide materials for evolution. TF gene loss leads to slightly different results than the loss of functional genes. Loss of TFs decelerates the molecular evolution of TGs and increases the level of their expression (at least in mice) (Figs. 2 and 3). The evolutionary rate of a protein is strongly negatively correlated with its expression level and the strength of any functional constraints (53–55). The loss of TFs in mammals enhances, rather than reduces, functional constraints on their TGs. A positive correlation exists between the loss of TFs and mammalian ancestral traits (Fig. 4). Compared with humans, the mouse niche may be closer to that of mammalian ancestors, with mice possibly able to more easily retain TG functional constraints after loss of TFs.

Many TFs have been lost during mammalian history, especially around the K–Pg boundary or afterwards. The massive extinction of dinosaurs as predators may have reduced functional constraints on relevant transcription machineries in our ancestors. As a result, TFs were lost unless new species shifted their habitats into recently opened niches and developed new lifestyles adapted to the new environments. However, TF loss has enhanced functional constraints on any TF TGs surviving to the present day. Mammals that lost these TFs may have increased their survival rate by retaining the ancestral traits in an unchanged niche. When an old niche was occupied, a less-adapted species with a less-altered or insufficiently altered transcriptional machinery may have needed to take risks to adapt to a new niche and change traits for survival. In this study, we have uncovered two main findings. First, the number of TFs increased greatly during the early stage of mammalian species formation; later, however, TF losses predominated during the course of macroevolution. Second, in the face of serious environmental changes, such as the K–Pg boundary, the loss of such TFs rewired the regulatory network, set functional constraints on TGs, and facilitated organismal survival.

## DATA AVAILABILITY

TF data, phylogenetic trees, and other data related to our research: (https://drive.google.com/drive/folders/1dN3Tav55yMuHxlVbLOYsPFPHv6I6C9Ls?usp=sharing) …

## ACKNOWLEDGMENTS

We express our deep gratitude to Z. Zhang, W. Sun, Q.-Y. Yu, W. Wei, H.-B. Zhang and Y. Takahiro for their valuable comments and suggestions on the manuscript. We thank B. Goodson, Edanz Group, for editing the English text of a draft of this manuscript.

## FUNDING

This study was supported by a Grant-in-Aid for Scientific Research (B) (19H04070) and a Grant-in-Aid for JSPS Research Fellows (18F18385) from the Japan Society for the Promotion of Science, a University of Tokyo Grant for PhD Research, and funding from the China Scholarship Council (CSC).

## CONFLICT OF INTEREST

The authors declare no conflict of interest.

## SUPPLEMENTARY DATA

Supplementary Data are available at NAR online.

**Fig. S1.**
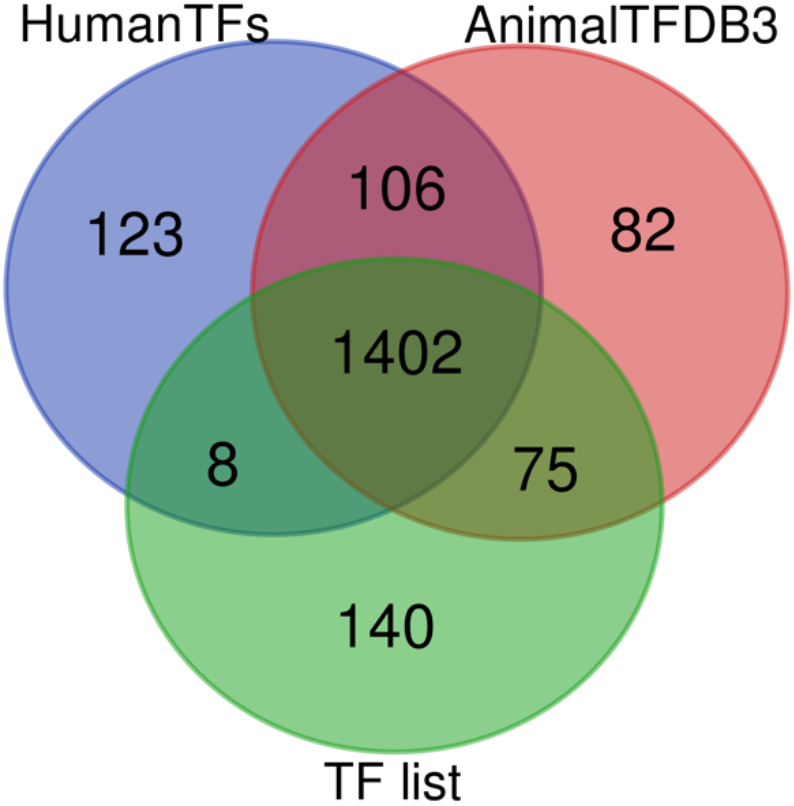
Venn diagram of human transcription factors (TFs) listed in HumanTFs, AnimalTFDB3 and our database. Numbers in blue, red and green circles are the number of human TFs in HumanTFs, AnimalTFDB3 and our database, respectively. Numbers in overlapping regions are the number of TFs shared by the different databases.

**Fig. S2.**
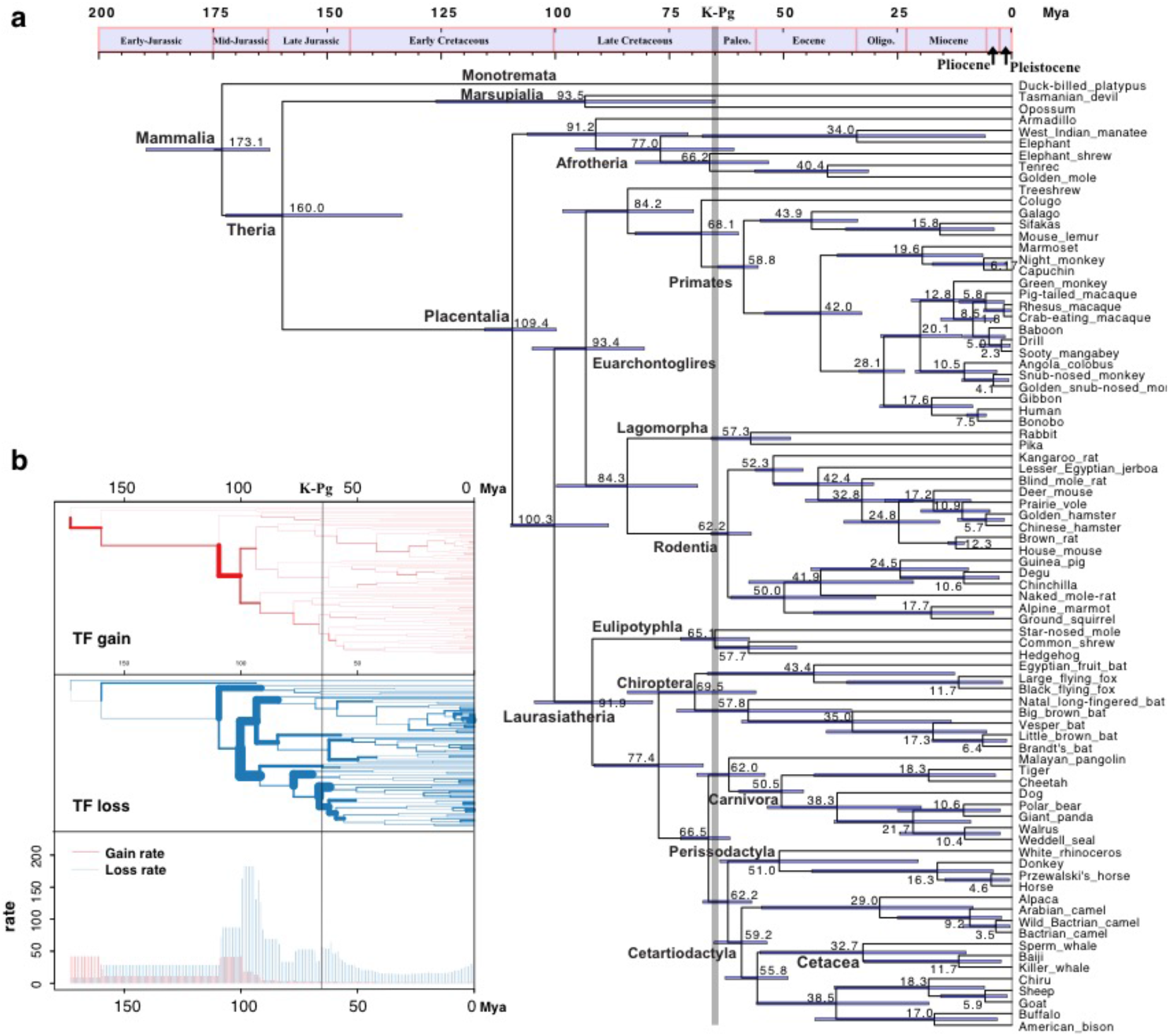
Simplification of transcription factors (TFs) during mammalian evolution. (A) Time tree of 82 mammals based on protein sequences. Numbers at internal nodes are estimated divergence times (Mya), with 95% credible intervals indicated by horizontal bars spanning nodes. (B) Atlas of TF gain and loss during mammalian history. The width of each branch is proportional to the rate of TF gain (or loss).

**Fig. S3.**
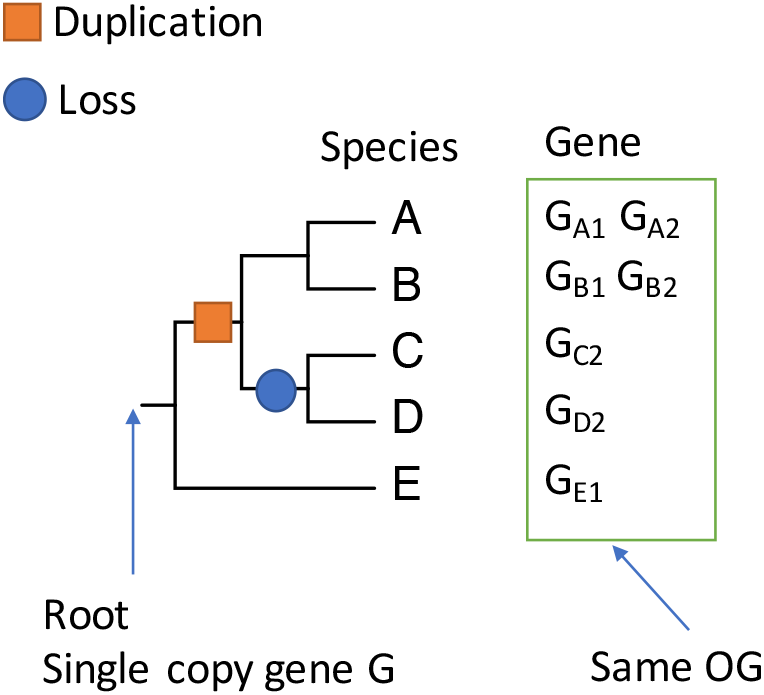
A simple example of an orthologous group. The orange square indicates a duplication event, and the blue circle represents a loss. The genes in the green box belong to the same OG and all originated from the single-copy gene G at the root.

**Fig. S4.**
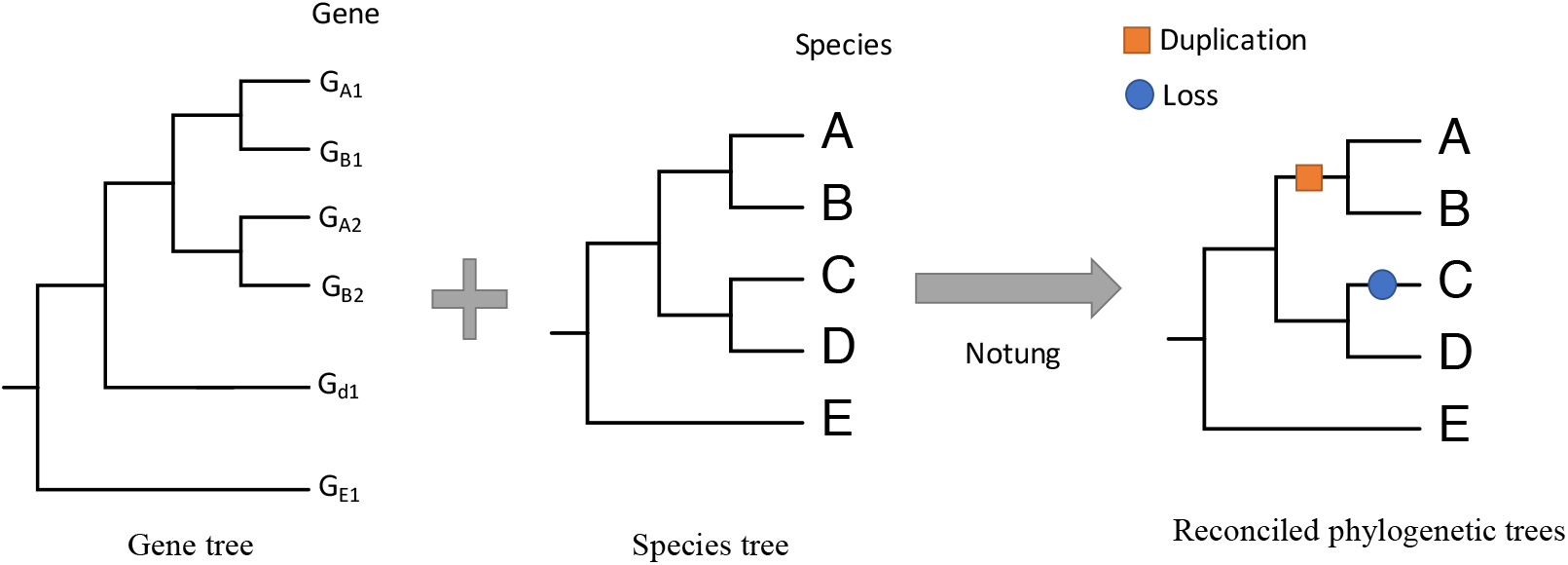
Inference of duplication and loss by gene-tree–species-tree reconciliation. G, gene. A–E, species. The orange square indicates a duplication event, and the blue circle represents a loss. Lines are branches.

**Fig. S5.**
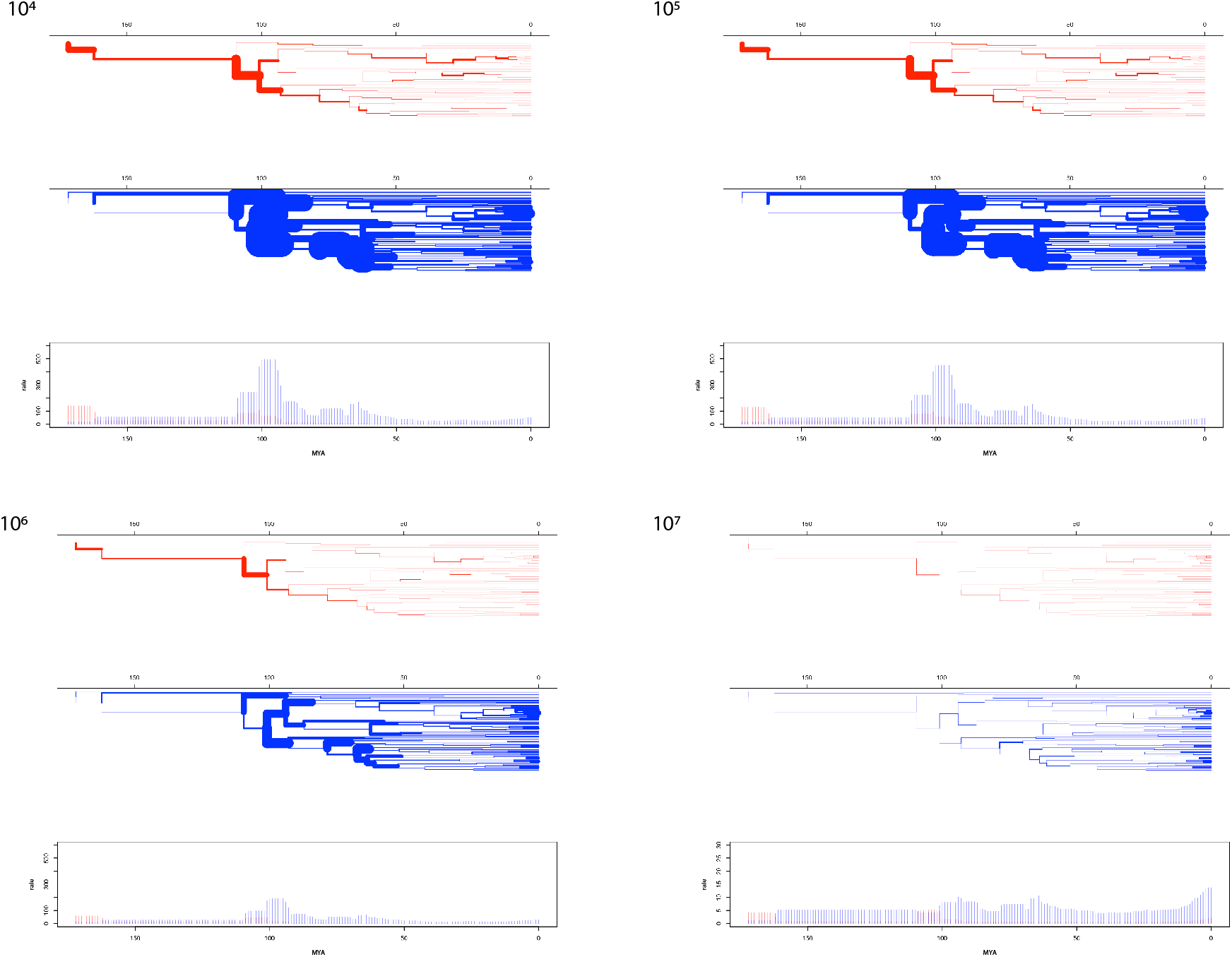
Transcription factor (TF) events inferred using four different effective population sizes—10^4^, 10^5^, 10^6^ and 10^7^—as a threshold when considering incomplete lineage sorting. TF losses and gains are represented by blue and red, respectively. The width of each branch is proportional to the rate of TF gain (or loss).

**Fig. S6.**
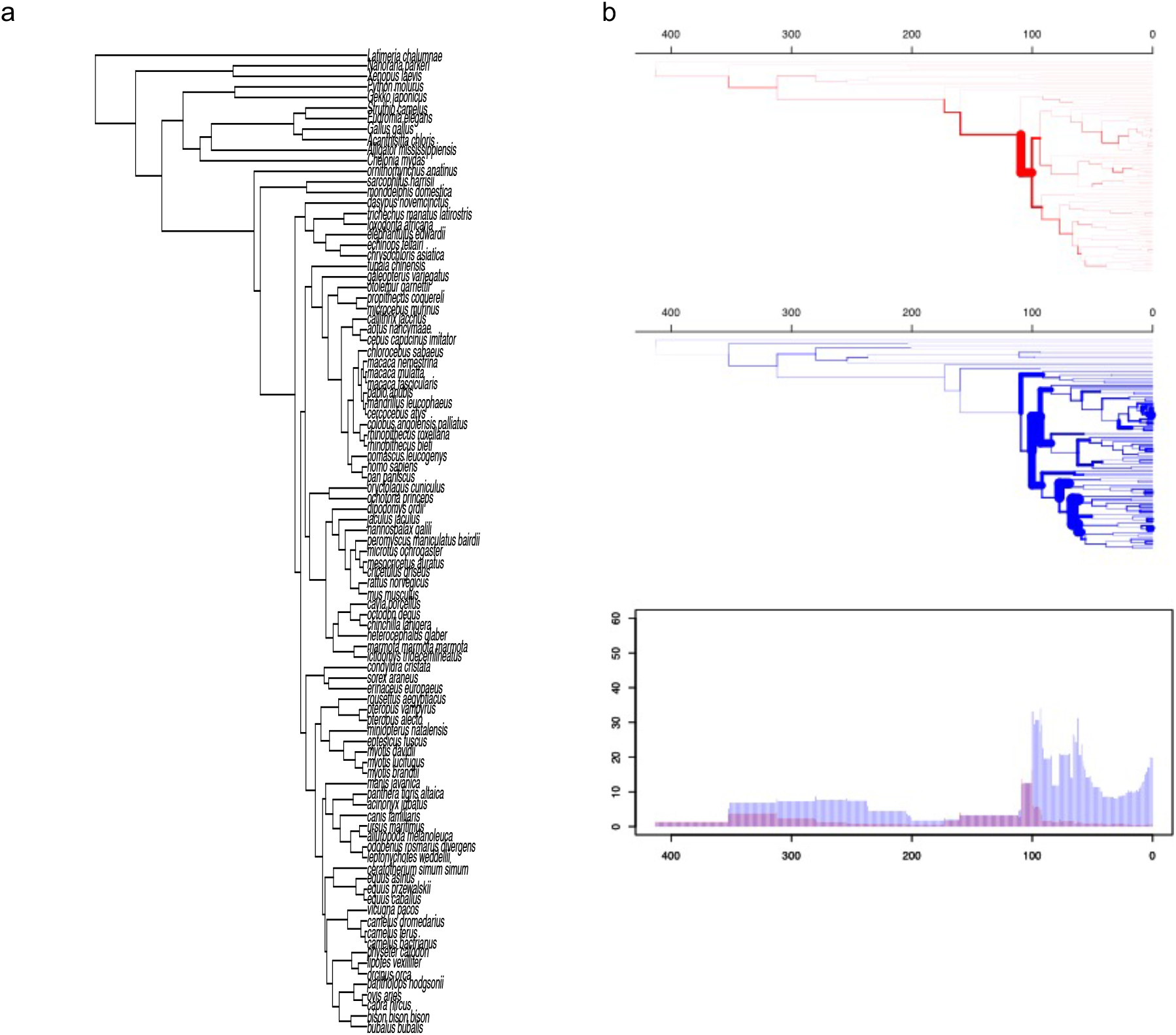
Transcription factor (TF) events in Atlantogenata inferred using an effective population size of 10^6^ with outgroup species included. TF losses and gains are represented by blue and red, respectively. The width of each branch is proportional to the rate of TF gain (or loss). Divergence times of outgroups are from Timetree (30).

**Fig. S7.**
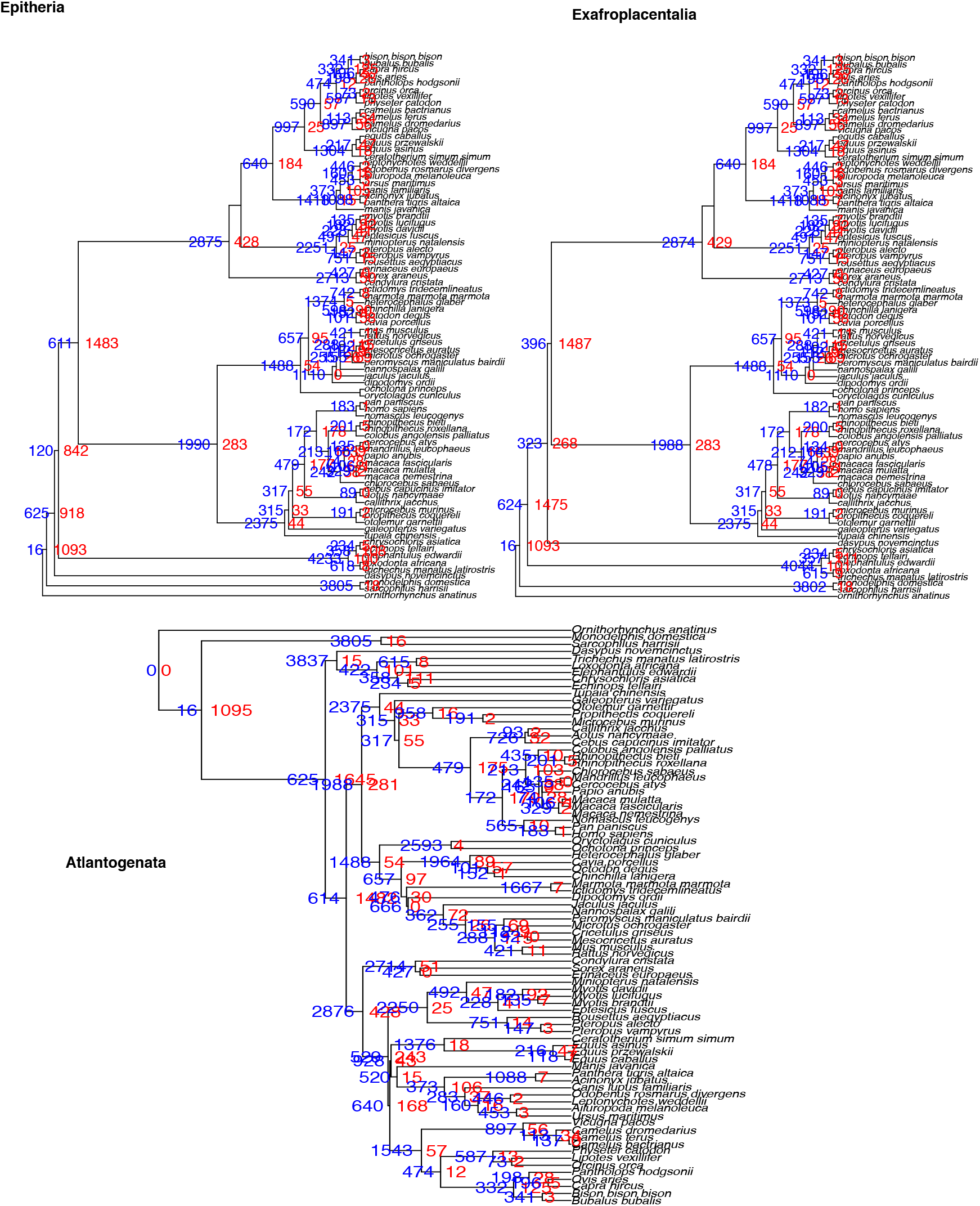
Transcription factor (TF) events in Atlantogenata, Exafroplacentalia and Epitheria inferred using an effective population size of 10^6^. Numbers of TF loss and gain events are indicated in blue and red, respectively.

**Fig. S8.**
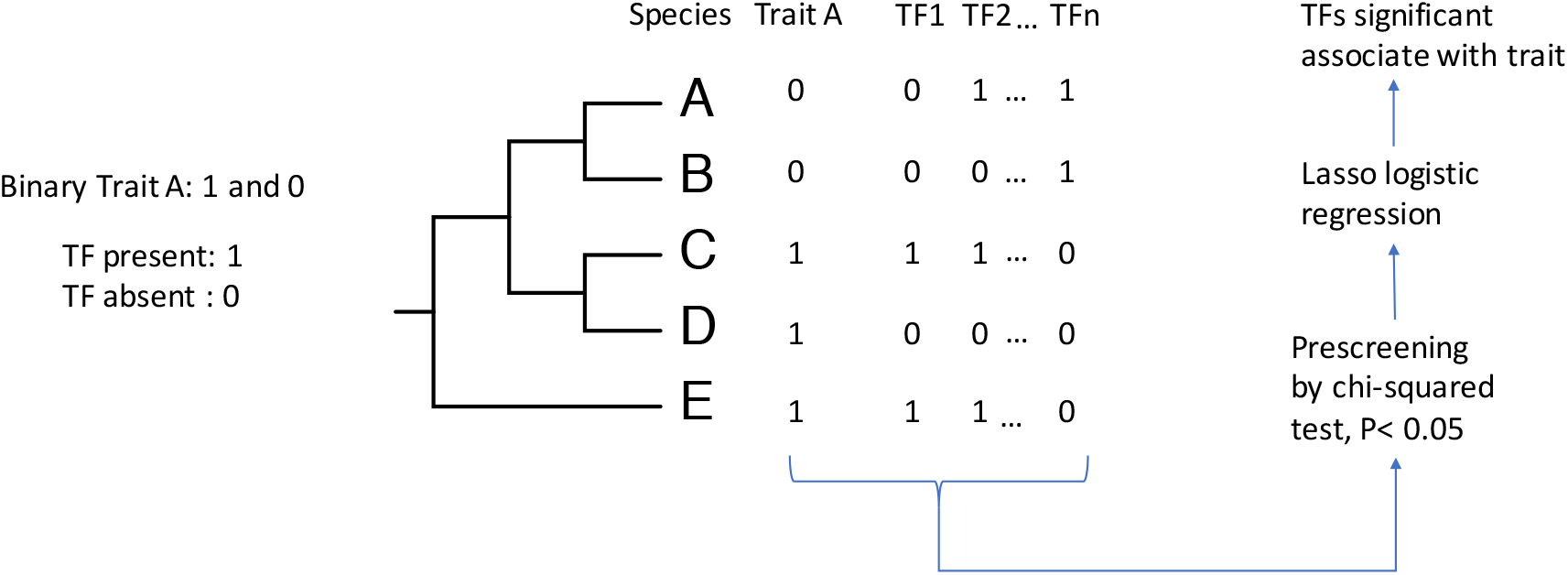
The pipeline used to detect an association between TF presence/absence and life history traits by lasso logistic regression.

(Supplementary Tables 1–3 are included as separate Excel files.)

Table S1. List of TFs with orthologous group annotations.

Table S2. Evolutionary rates of target genes experiencing TF loss over the long term.

Table S3. Association of TFs with mammalian traits based on lasso logistic regression.

## Notes

https://drive.google.com/drive/folders/1dN3Tav55yMuHxlVbLOYsPFPHv6I6C9Ls?usp=sharing

### Competing Interest Statement

The authors have declared no competing interest.

